# Estimating Wingbeat Frequency on Hummingbirds using a No-labeling Learning Computer Vision Approach

**DOI:** 10.1101/2024.10.04.616678

**Authors:** Maria Ximena Bastidas-Rodriguez, Ana Melisa Fernandes, María José Espejo Uribe, Diana Abaunza, Juan Sebastián Roncancio, Eduardo Aquiles Gutierrez Zamora, Cristian Flórez Pai, Ashley Smiley, Kristiina Hurme, Christopher J. Clark, Alejandro Rico-Guevara

## Abstract

Wingbeat frequency estimation is an important aspect for the study of avian flight, energetics, and behavioral patterns, among others. Hummingbirds, in particular, are ideal subjects to test a method for this estimation due to their fast wing motions and unique aerodynamics, which results from their ecological diversification, adaptation to high-altitude environments, and sexually selected displays. Traditionally, wingbeat frequency measurements have been done via “manual” image/sound processing. In this study, we present an automated method to detect, track, classify, and monitor hummingbirds in high-speed video footage, accurately estimating their wingbeat frequency using computer vision techniques and signal analysis. Our approach utilizes a zero-shot learning algorithm that eliminates the need for labeling during training. Results demonstrate that our method can produce automated wingbeat frequency estimations with minimal supervision, closely matching those performed by trained human observers. This comparison indicates that our method can, in some scenarios, achieve low or zero error compared to a human, making it a valuable tool for flight analysis. Automating video analysis can assist wingbeat frequency estimation by reducing processing time and, thus, lowering barriers to analyze biological data on fields such as aerodynamics, foraging behavior, and signaling.

## Introduction

Hummingbirds (family Trochilidae) are renowned for their agile flight and unique ability to perform sustained hovering among birds (e.g., Altshuler and Dudley 2002, Skandalis et al. 2017), which is a fundamental component of their ecological and social interactions (e.g., Warrick et al. 2009). This ability is granted by a combination of exceptional anatomical and physiological adaptations, most notably their rapid and precise wingbeats. Studying the flapping frequency of hummingbirds has implications in several fields, for example, robotics, aerospace engineering, and evolutionary biology (e.g., Altshuler et al. 2004, Tobalske et al. 2007a, Nan et al. 2019, Tu et al. 2020, Wilcox and Clark 2022, Zhao et al. 2022, Jensen et al. 2024)

Hummingbirds’ versatile flight allow them to perform complex maneuvers such as hovering, backward flight, and rapid turns (e.g., Martínez et al. 2015a, Dakin et al. 2018). These capabilities are crucial for their feeding behavior, enabling them to access nectar from pendant flowers with precision and efficiency (e.g., Stiles 1971, Inouye et al. 1991). However, the wingbeat frequencies off the majority of species remain unknown, mostly due to logistical limitations (e.g., capturing and processing high-speed video). Hummingbirds display minimal variation in wingbeat frequency across many conditions, including forward flight, hovering, and different flight speeds for instance their wingbeat frequency remains relatively constant during both hovering and forward flight (Chai et al., 1999; Tobalske et al., 2007b; Clark and Mistick, 2018), or even when facing aerodynamic challenges such as reduced airfoil area during molt (e.g., Chai 1997, Díaz-Salazar et al. 2023), therefore characterization of this flight parameter will be informative for a variety of contexts.

Greenewalt (1960) pointed out that, allometrically, hummingbirds actually have lower wingbeat frequencies than it would be expected for their size, for example, a 10-gram chickadee has a higher wingbeat frequency than a 10-gram hummingbird. Wilcox and Clark (2022) derived the expected allometric relationship for hummingbird wingbeat frequency, taking into account their wing length allometry. Compared to most birds, hummingbirds have longer wings relative to their body size. This wing length allometry endows them with high maneuverability and sustained hovering capabilities. Wilcox and Clark’s findings reveal that the allometric slope of wingbeat frequency in 30 species from 37 out of the bee hummingbird clade closely matches the predicted values, highlighting their unique wing structure and wingbeat frequency play a significant role in their flight mechanics and presents a compelling area for further research (Greenewalt, 1960; Wilcox and Clark, 2022). However, data on a larger proportion of the over 350 species of hummingbirds, across their large elevational and latitudinal ranges is needed to test the robustness of that and other morphofunctional flight patterns.

Currently, wingbeat frequency estimations have been performed visually by digitizing landmarks frame by frame and counting wing beats in high-speed videos, or from processing of sound recordings of hovering (e.g., Steen 2014; Sotavalta 1952). Automating video analysis could streamline post-processing, alleviating the need of significant amount of time and effort. However, there are several common limitations of automating data collection from videos. First, excess videos must be acquired to train the automated program in addition to what is needed to investigate the question of interest. Second, illumination, video frame rate, framing, focus, lighting, orientation of the hummingbird, and occlusion by other objects in the frame make it difficult to determine specific motion patterns and can affect data quality. Third, in supervised machine learning, these datasets often need to be annotated by a human expert (e.g., labeling and manually bounding boxing the study subjects in the video frames). These annotations, typically done frame by frame, can be time-consuming, and arduous to perform.

In this scenario, recent zero-shot learning methods, a machine learning approach where the model is capable of analyzing data it has never seen before (Xian et al., 2019), are particularly appealing. Zero-shot learning uses pre-trained models (available online, e.g., YOLO-NAS model 2023, OPEN CLIP Model 2024), which are trained in different tasks (e.g. object detection/classification, Supplementary Information S1.1), and benchmark datasets that can be adapted to new and unknown tasks. This procedure could work for particular research questions, even though the models are not specifically trained to solve a given problem, they still can generate accurate results. Nguyen et al. 2024, uses this technique to detect and segment the pixels where animals are present on the image, such as dolphins, frogs, and birds, among others, with the particularity that the ambient texture of the background and colors tend to mix with the ones from the animals; their study uses a deep learning image feature extraction method and compare them with the cosine similarity. In terms of bounding box image detection the YOLO (You Only Look Once) family detectors—widely recognized in computer vision (see Jiang et al. 2022)— have been used by different researchers (e.g., Ma and Yang 2022) modified the YOLOv5 net for animal detection).

The versatility of OPEN AI CLIP algorithm (Radford et al., 2021) allowed improvement in zero-shot algorithms. Shin et al 2023 used this method to create a non-supervised segmentation algorithm; which has also been used for the recategorization of birds with only a description of color and textural features (Wu and Maji, 2022). Computer vision approaches have also been used for wingbeat frequency analysis, for example segmenting and tracking infrared videos of flying bats to evaluate their wing beat position relative to their body (Breslav et al., 2012). Also, Ling et al. 2018, used three-dimensional analysis by capturing stereo images of jackdaws and rooks to track motion measuring velocities, accelerations along with wingbeat frequency and claim that these calculations help to understand the flight kinematics of birds.

For hummingbirds, the studies in computer vision have focused on the classification of frames in a video identifying hummingbirds in their nests (Serrano et al., 2018); or in detectors as the one proposed by Weinstein (2015, 2018), who compared each video frame of an image with a background image to determine the presence of a new object to filter videos in flowers (an alternative to camera traps and triggering systems, (e.g., Rico-Guevara and Mickley 2017). Martínez and collaborators 2015b used dense optical flow features to characterize the hummingbird wings motion measuring the global angular acceleration of the wing and the global wing deformation. Hummingbird flight has inspired studies in the development of flapping wing Micro Air Vehicles (MAVs e.g., Nan et al. 2019, Tu et al. 2020). Lastly, Zhao and collaborators (Zhao et al., 2022) designed an optimization algorithm for engineering processes through hummingbird flight simulation.

Given the multidisciplinary interest on this topic, and the absence of studies focusing on methods to study flight characteristics efficiently across species, we focused on developing an automated system for detecting, classifying, tracking, and quantifying wingbeat frequency in hummingbirds while hovering. We evaluate our approach using videos of free-living hummingbirds without any illumination or scene control, which permits the processing of videos collected under a variety of conditions matching the varied environments where hummingbirds live.

## Methods

Given our goal of developing a system able to detect, and re-identify objects (specified by the user employing prompt texts), track them, and analyze their movement along the video, we employed a Zero-shot approach that allows use under different scenarios, and does not only detect hummingbirds but can be easily adaptable to other animals or flying objects (e.g., MAVs). This computer vision algorithm to study hummingbird flight consists of four specific modules (Figure 1):

**Figure 1.**
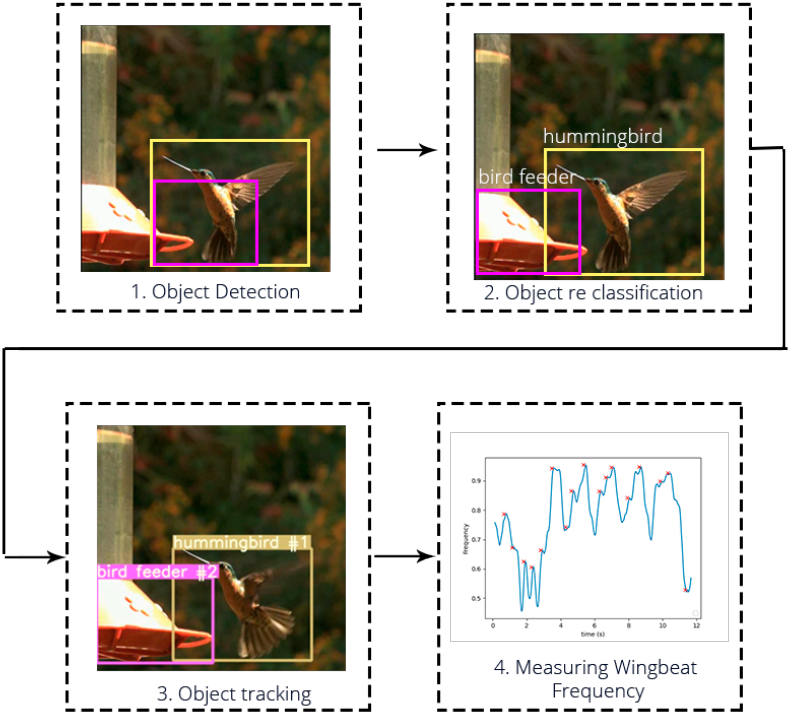
Block diagram of the proposed algorithm. 1. An object detection stage using YOLO-NAS; 2. A classification stage using text-prompt with OpenAI Clip; 3. An object tracking stage; and 4.The proposed Wingbeat Frequency Estimation method.

1. A detection module using a YOLO-NAS algorithm (Aharon et al., 2023) that allows us to obtain objects in the image by localizing and enclosing them into a bounding box.
2. A re-classification method by using the OpenAI Clip algorithm (Radford et al., 2021) and a text prompt of the classes of interest, e.g., hummingbird, flowers, etc.
3. A tracking system, by comparing image features, obtained by the Open Clip algorithm, from sequential frames of the obtained bounding boxes from the detection module.
4. A frequency analysis that allowed us to approximate the count of the wingbeats of a focal hummingbird from the detection boxes across video frames.

### Dataset and Computational Resources

We recorded high-speed videos of hummingbirds performing static hovering and visually measured their wing flapping frequency (e.g., Steen 2014). The videos were taken with fixed point cameras (Fastec HS7, a Fastec TS5, and six Chronos 2.1) using a minimum resolution of 800 × 600 pixels and recorded at 240 and 500 frames per second. The RGB color space of each frame is analyzed individually through the proposed pipeline. The illumination conditions as well as the devices were not standardized, however, we selected video sequences where the hummingbird was on a profile view and was the main object on the scene, without backlighting or illumination changes. A total of 14 videos were selected to compare the performance of the proposed algorithm with the visual estimation method, ensuring that each video contained only one bird flying in the shot (although other birds could be statically present on the feeders). This selection facilitated the analysis by focusing on a focal bird as the main object in the scene. We counted the wingbeats on videos between 300 to 1000 frames of duration, having around 21 to 90 wingbeats per video.

### Object detection method

The first step consists of detecting the objects in the image and separating their pixels from the rest of the image (Figure 2) Detection Module. The detection locates an object and encloses it into a bounding box that contains every pixel of interest. A robust algorithm should be agnostic to the pose, illumination, or deformation of a given object. One of the most popular algorithms for this task are the ones denominated as YOLO Family (Jiang et al., 2022). These algorithms used a deep learning architecture trained in a two-task optimization function for detection and classification (See Supplementary Information S1.1). The generalization ability of these methods allows them to use pre-trained models and adapt them to specific tasks which makes them perfect for zero-shot learning analysis. Also, it is characterized by being one of the fastest and smallest methods for this kind of problem so it makes them suitable for video analysis. For this study, we apply the YOLO-NAS architecture (Aharon et al., 2023). It uses a neural architecture search technology, which means that the architecture was automatically designed to improve the quantization and accuracy of the models and optimize the hyperparameter search. By doing a visual analysis, we determined that the YOLO-NAS algorithm was better suited for the job and we corroborated it by analyzing the frame detection on the videos. Finally, it is worth mentioning that the output of the detector algorithm could predict multiple boxes for the same object, and to determine the best prediction an algorithm called non-maximum suppression is used to discard those that overpass a certain overlap threshold.

**Figure 2.**
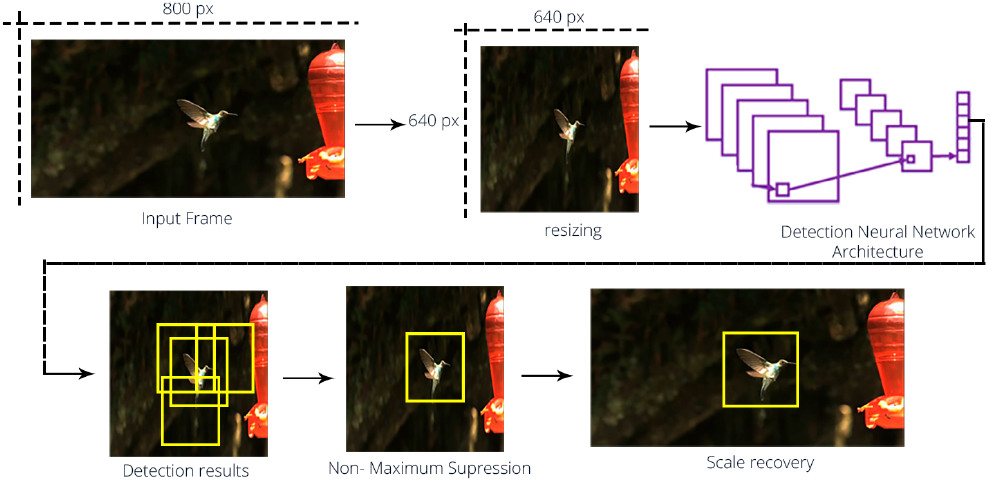
Overlook of the detection process. First step consists of resizing the image to 640 px x 640 px, the algorithm returns the boxes where the objects present on the scene are located. Several boxes could be detected for each object, then the non-maximum suppression algorithm is used and the most suited box is selected. Finally the image scale for the box is restored.

As the proposed algorithm is a zero-shot learning algorithm for both detection approaches we applied pre-trained models used for inference. These pre-trained models were originally trained on the COCO dataset (Lin et al., 2014). These datasets contain 80 object classes with almost 330.000 labeled images with the correspondent bounding boxes ground truths. Among those classes, class 14 is defined as hummingbird; which means that the algorithm trained on this dataset would take hummingbird features into account. The classification was not considered as these algorithms tend to have less precision in this task than other methods and a re-classification is performed in a followup module. By doing so, we were able to use more robust and specific algorithms for the classification and tracking not only of the hummingbird but also other elements given by the user with text prompts (see: Classification Module).

### Classification Module

As mentioned, the algorithms from the YOLO Family are robust in the detection object part but tend to have a minor performance during the classification task without fine-tuning the weights. For the proposed algorithm, we use a re-classification module (Figure 3) which is based on the Large Scale Learning Model (LLMs) CLIP (Radford et al., 2021). These models are trained on an extensive dataset that maps the text features to the image features through the minimization of the distance between them, normally the cosine distance (See Equation 1), searching for the similarity between the bimodal (text and image) inputs. Thus, it is feasible to perform a classification of the objects without performing a re-training and limiting the search to the objects of interest.

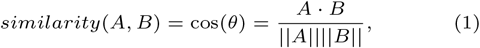

where,

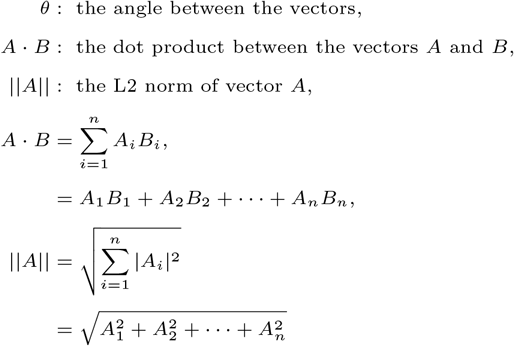

**Figure 3.**
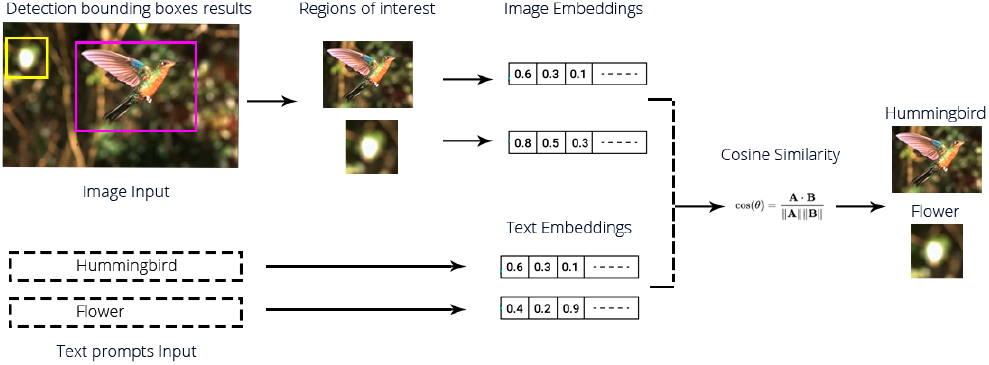
Overlook of the classification process. We are using a contrastive learning tool where feature embeddings of the detected regions on the image and the given vocabulary are contrasted, by using the cosine similarity, to obtain the final classification

We use the approach for two main functions :

- First, we obtained image features from the region of interest within the detected bounding boxes and text features from the text prompts given in advance. For our experiments, we used tow words: hummingbird and flower. However, this could be easily changed in the code to be adapted to any user needs. The process to perform the reclassification is as follows:

1. Obtain text features from the given classes.
2. Obtain image patches using the detected bounding boxes for the frame to analyze.
3. Obtain image features for each of the extracted patches.
4. Calculate the cosine distance between each image patch feature and each text prompt.
5. Select the minimum distance to assign the analyzed object to a particular class.
6. Repeat the process on each of the video frames.

- Second, we transferred the image features obtained from each region of interest to the tracking module (See subsection Tracking module) to compare the detected objects on sequential frames.

### Tracking module

This module uses the Deep SORT algorithm (Wojke et al., 2017), which receives the extracted image features from the Classification Module and the detection boxes from the detection task (Object detection method). In the first video frame, these features are stored, and each detected object is assigned a tracking ID based on the class name and a sequential number (e.g., “hummingbird 1”). Each detected object is represented by a feature vector. In subsequent frames, comparisons are made between the stored data from previous frames and the current frame. A Kalman filter predicts the next position of each object, and the Euclidean distance between the predicted box and the detected box in the current frame is calculated. Additionally, the features of each object are used to distinguish between objects that may have similar positions by calculating the cosine similarity (see Equation 1) between the image feature vectors. If the similarity is below a given threshold, the new features are assigned to the matched track ID; otherwise, a new track is created, and a new set of tracking IDs is generated.

### Measuring wingbeat frequency

In this research, we used a similar approach to the one used by Zhang et al. 2009, where they performed an automatic frequency analysis of bird’s flight by using a detection and tracking algorithm, they remark that the area of the detection box would be wider when the bird performs an upstroke (upward stroke of the wing) while it would reduce when the downstroke occurs. This assumption is not entirely true when it comes to analyzing hummingbirds. Due to its velocity and flexibility, the upstroke of a hummingbird could easily have a similar area to the surrounding detection box of the downstroke (See Supplementary Information S1.3). For this reason, we proposed to use the Jaccard index or intersection over union IoU (See Equation 2) of boxes of consecutive frames. The idea is to measure the location change of the box on consecutive frames, and by doing so, measure the frequency of these changes.

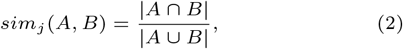

Then, we use the short-term Fourier transform (STFT) to monitor the distribution of the frequency during a time slap of the generated signal (Zhang et al., 2009). This frequency-time analysis is a collection of multiple discrete Fourier transforms from regular overlapped windowed intervals of the signal and it is represented by the function given in Equation 3 (Kehtarnavaz, 2008).

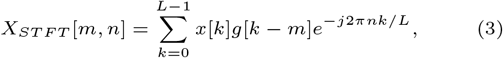

where,

*x*[*k*] :Corresponds to the signal,

*g*[*k*] :Denotes the analysis window,

*L* : Denotes the length of *g[k]*

Finally, we get the high-frequency signal of the spectrum, apply a Savitzky-Golay filter to smooth it, avoid small movements of the IoU to be counted as flaps, and find the peaks of the resulting signal (Figure 4), shows the proposed measuring method. The final signal with the peaks counted as upstrokes of the hummingbird flight.

**Figure 4.**
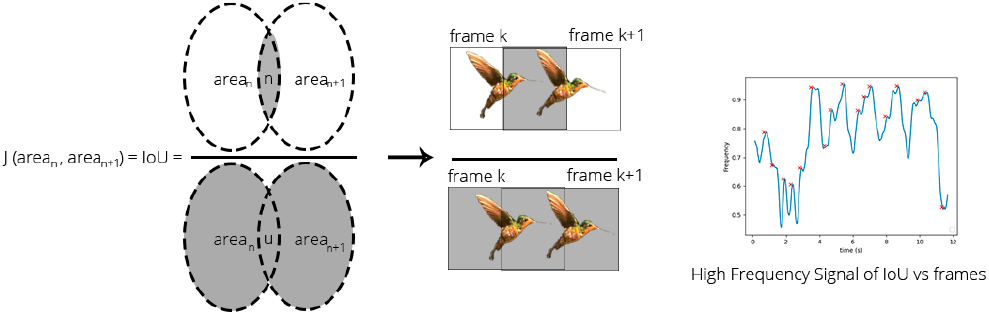
The proposed wingbeat measuring method. By using the concept of the Jaccard’s coefficient, we analyze consecutive frames by obtaining the bounding boxes area intersection over the bounding boxes area union. Giving a measure of movement of the hummingbird

To compare the results we counted the peaks for each of the analyzed videos and an expert biologist counted the wingbeats on each video. Then, we transform this count to flight frequency by the formula presented in Equation 4.

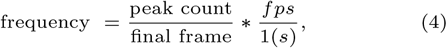

And we measure the error by using the percent error given by Equation 5

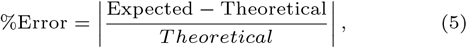

As the proposed algorithm is divided into three main steps (detection, classification, and wingbeat frequency measuring) we evaluate each step separately, focusing only on the class of interest: hummingbird. For each step, we used three common metrics on the computer vision field named: Precision, Recall, and F1-score, which depend on the True Positive (TP), False Positive (FP), and False Negative (FN) values.

- Precision: Evaluate the model by giving a proportion of the true positives among all the positives given by the model. Its results are interpretable as percentage values (See Equation 6).

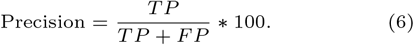
- Recall: Evaluate the model by giving a proportion of the true positives among all the real labeled positives or ground truth. Its results are interpretable as percentage values (See Equation 7).

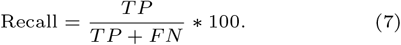
- F1-score: This measure combines Precision and Recall by calculating the harmonic mean (See Equation 8).

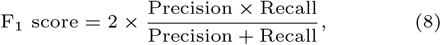

where, 1.0 is the best possible value.

## Results

### Detection

To validate that the YOLO-Nas detection algorithm performs correctly in the selected dataset, we obtained 20 frames per video, randomly chosen, and manually labeled them, obtaining the bounding boxes where hummingbirds appear. In this way, a total of 198 frames were analyzed, in some of the frames there was more than one hummingbird on the scene (one flying and the rest on the bird feeder), generating a sum of 222 detections. Then the intersection over union (IoU) metric was calculated between the labeled bounding box and the obtained with the model. Supplementary Information (S1.2) shows the distribution of the IoU among the predictions, it is possible to observe that most of the detections have an IoU greater than 0.5 which means that the majority of the detections were done correctly. In the context of detection, a true positive value is given when the IoU is greater than a given threshold (normally the threshold is a number between 0.5 to 0.95), in our case the threshold chosen was 0.5; a false positive is given when the IoU is lower than the threshold; and the false negative value is given when there was not a box that detected the object of interest. A true negative value will be every frame where an object was not labeled and not detected, this value does not have interest in this analysis. In our experiment, *TP = 180, FP = 74* and *FN = 39*, which achieves a *Precision = 71 %, Recall = 82 %* and *F1score = 0*.*76*. These results confirm that the YOLO-Nas algorithm is suited for the detection of different species of hummingbirds.

### Classification

For the classification algorithm, we focused on the classification of the class: “hummingbird”. We did not count the classification by frame, as in the evaluation of the detection algorithm, instead, we took into account the interest class by video. This means that if an object is classified (or misclassified) as a hummingbird, it will count as one until the video is finished or it has a change of class. Then, the same object could be considered as a TP and FN (or TN and FP) during the same video. In this way, we can obtain an insight into the full video (Supplementary Information S2 Figure. S3 and Table. S1).

Summing up the results, we obtain *TP = 16, FP = 0, FN = 3* and *TN = 8*, which achieves a *Precision = 100 %, Recall = 84 %* and *F1score = 0*.*91*.

### Measuring wingbeat frequency

The method proposed counts the picks detected on the high-frequency component of the IoU signal (Supplementary Information S3, Table. S2.). Even though the results are affected by the errors made by the detection, classification, and tracking algorithms, in most of the analyzed videos we obtained values similar to the manual counts (Supplementary Information S3, Table. S2., Figure 5). Supplementary Information S3, Figure. S4., shows two frequency vs. time plots, where it is possible to corroborate the pick count done by the algorithm with the proposed method. For two of the videos, it was not possible to count the peaks as the errors with the detection algorithms were too high (the birds detection was on and off), so the consecutive frames IoU metric were affected.

**Figure 5.**
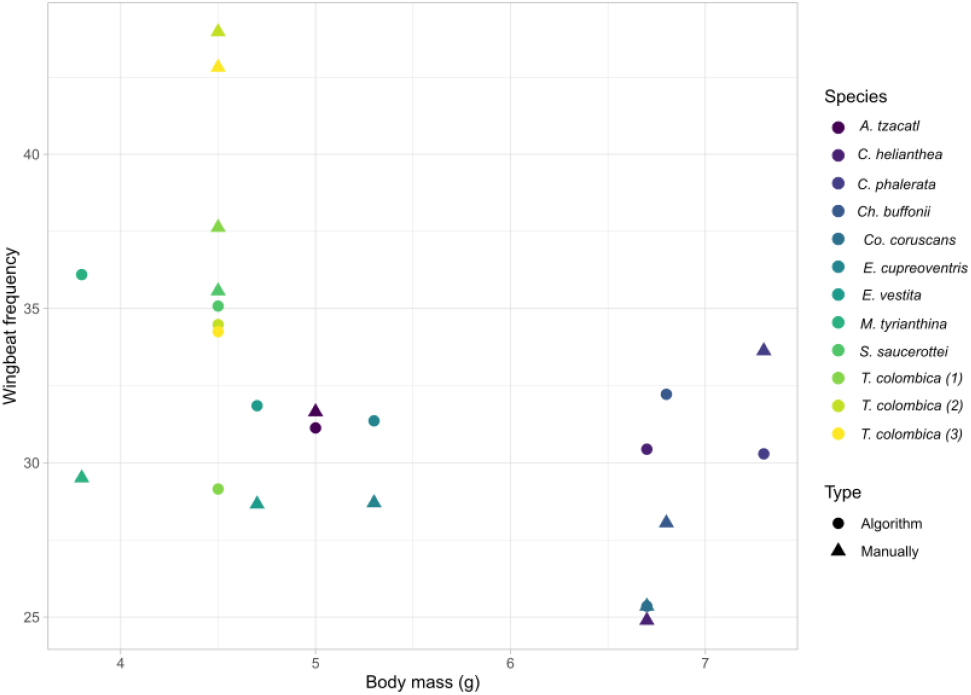
Wingbeat frequency and body mass (from Ayerbe-Quiñones (2024)) for the 14 individuals of 12 hummingbird species. U for unknown sex of individuals. Filled circles denote wingbeat frequency measured manually and filled triangles wingbeat frequency measured by the algorithm. The full scientific names and sex of each individual is provided in Supplementary Information (S3), Table S2.

## Discussion

Wingbeat frequency and stroke amplitude are among the most straightforward flight parameters to measure, making them ideal starting points for researchers conducting comparative studies in basic kinematics. These parameters are often the first to be modulated during different types of flight, such as turning or executing aerial maneuvers, providing valuable insights into the fundamental mechanics of flight (Chai et al., 1999; Tobalske et al., 2003; Altshuler et al., 2010). In hummingbirds, these kinematic traits are not only crucial for understanding flight mechanics but are also linked to specific behaviors, such as courtship displays (Feo and Clark, 2010). Our study presents a method that successfully approximates the wingbeat frequency for the analyzed subjects. The approach comprises four main stages and operates as a zero-shot learning technique, requiring no retraining. This allowed to validate the algorithm without the need to recollect training data, saving both computational and labeling (observer time) resources.

The detection algorithm achieved an *F1-score* of 0.76 and a *Recall* of 82 %, indicating that it detected nearly every hummingbird identified by human observers, with the main challenges arising from filming issues such as blurriness and partial occlusion of the bird. The classification algorithm performed an *F1-score* of 0.91 reflecting low rates of false positives (FP) and false negatives (FN), and correctly distinguishing hummingbirds from other objects in the scene. Both algorithms proved robust across different hummingbird species and minor variations in flight and positioning. We present estimated wingbeat frequencies in relation to the reported body mass for the species for both the manual and the algorithm counts to easily visualize their differences (Figure 5). We found a reduction in flapping frequency with increased body mass, adding new species to a trend previously reported (Altshuler and Dudley, 2003; Groom et al., 2017; Krishnan et al., 2022). A higher wingbeat frequency in smaller species has been proposed to be linked to shorter wings being able to move through the air and complete wingbeat cycles faster (Hunter and Picman, 2005; Steen et al., 2020).

We did not include in the comparison graph across species a 30-second video of *Ensifera ensifera* (Supplementary Information S3, Figure. S5.), which illustrates both the strengths and limitations of the tool. In the initial segment, where the bird is viewed from the back, the detection algorithm struggles, leading to missed detection in some frames. The classification algorithm also misidentifies the bird in this view. However, when the bird turns to a profile view, both the detection and classification algorithms perform significantly better, and the accuracy of wingbeat counting improves, with human experts counting 32 wingbeats compared to 31 by the algorithm. In the final segment, the bird’s rapid change of direction causes the tracking algorithm to fail. It’s important to note that the system relies on three distinct machine learning models, none of which were trained on the data analyzed in this study. To improve these results, fine-tuning the detection and classification algorithms could be a valuable step. Furthermore, while we collected videos in the wild, the algorithm performed best when the subject occupied a significant portion of the frame and the shot was taken from a profile angle (Supplementary Information S1.3).

The proposed algorithm offers an automated, easy-to-implement method for counting wingbeats in videos, enabling data collection by non-experts in the field, such as using high-speed cameras near tourist spots with feeders. For those seeking a more accessible approach, you can also use your smartphone to record the wing hum, then open the recording in free software such as Raven Lite (Charif et al., 2006). By setting the spectrogram parameters to a wide window, and look for and measure the dark band near 50 Hz to process the video and determine the wingbeat frequency with this software or manually. Our results show that it is possible to approximate wingbeat counts with a small margin of error. Of the analyzed videos, twelve had an error below 25 % in the estimated wingbeat frequency, five of them are under the 10 % error, and one video has an error above the 50 % (*Ensifera*). Upon closer examination of the videos with high wingbeat counting errors (See Supplementary Information S3, Figure. S5.), it became evident that these errors likely resulted from the hummingbird being too small relative to the frame, the video being too blurry for accurate analysis, or the bird position is not on a profile angle. These issues could be easily addressed by improving filming techniques to meet the minimum standards our analysis (Supplementary Information S4) helps to identify. In contrast, Supplementary Information (S3), Figure. S6. presents frames from the two videos with near to 0 % error in frequency. In these videos, the hummingbirds were minimal changes in the birds’ flight direction or orientation during analysis.

The algorithm faced challenges when processing out-of-focus videos, an issue particularly relevant for biologists, as the rapid movements of hummingbirds often cause cameras to lose focus. From a computer vision perspective, analyzing blurry images is inherently difficult because of the loss of high-frequency information. Additionally, our use of a zero-shot learning approach, which did not involve labeling any of the training images or videos, further complicated the analysis. To address these limitations in future work, we could train the detection and classification algorithms to enhance feature tracking, which could also be adapted to handle the different orientations of the bird in various video shots. In tests with videos where the hummingbirds remained in profile but at a different inclination, the algorithm performed well. However, when the bird was not in a profile view, the detection and classification algorithms began to fail. This issue could be mitigated in future studies by training these algorithms with sufficient labeled data, although this process is time-consuming. Automating the analysis of flapping frequency in videos will significantly enhance our understanding of this critical flight parameter across hummingbirds. However, capturing videos with ideal backgrounds and lighting conditions can be challenging in natural settings.

## Data Availability

Code is available at http://github.com/mxbastidasr/bird_zero_shot_tracking

## Author contributions

Conceptualization: M.X.B.-R., A.R.-G Video collection: M.J.E.U., D.A., J.S.R., E.A.G.Z., C.F.P. Data analysis: M.X.B.-R., A.M.F., M.J.E.U., D.A. Writing first draft: M.X.B.-R., A.M.F., A.R.-G Critical revision and editing: M.X.B.-R., A.M.F., A.S., K.H., C.J.C., J.S.R, E.A.G.Z, A.R.-G.

## Acknowledgment

The authors thank the researchers of Colibrí Gorriazul Research Station for their help in recording and processing videos. The members of the Behavioral Ecophysics Lab of the University of Washington for their insights on the manuscript. And finally, to Lucero Rojas and Parmenio Simbaqueba for their help in the maintenance of feeders and work at Colibrí Gorriazul Research Station and to Victoria Lizarralde for the maintenance of the feeders of Observatorio de Colibríes.

## Ethics

All work was done under Washington University ethics approval 4498-05 and Colombia Regional Environmental Corporation (CAR) permit number DJUR 50227001473.

