## Supplementary material for "Estimating Wingbeat Frequency on Hummingbirds using a No-labeling Learning Computer Vision Approach"

**S. Eq. 1 Detection loss function:**

$$L_{loss} = L_{bbox} + L_{bbox_{score}} + L_{class_{score}}$$

Where:

- $L_{bbox}$: Bounding boxes loss function.

$$L_{bbox} = \lambda_{coord} \sum_{i=0}^{S^2} \sum_{j=0}^{B} 1_{ij}^{obj}[(x_i - \hat{x}_i)^2 + (y_i - \hat{y}_i)^2] +$$
$$\lambda_{coord} \sum_{i=0}^{S^2} \sum_{j=0}^{B} 1_{ij}^{obj}[(\sqrt{w_i} - \sqrt{\hat{w}_i})^2 + (\sqrt{h_i} - \sqrt{\hat{h}_i})^2]$$

- $L_{bbox_{score}}$: Bounding boxes score loss function.

$$L_{bbox_{score}} = \sum_{i=0}^{S^2} \sum_{j=0}^{B} 1_{ij}^{obj} (C_i - \hat{C}_i) + \lambda_{noobj} \sum_{i=0}^{S^2} \sum_{j=0}^{B} 1_{ij}^{noobj}(C_i - \hat{C}_i)$$

- $L_{class_{score}}$: Classification loss function.

$$L_{class_{score}} = \sum_{i=0}^{S^2} 1_{ij}^{noobj} \sum_{C \in classes} (\rho_i(C) - \hat{\rho}_i(C))^2$$

Where:

- $1_{ij}^{obj}$ : $it\ is\ 1\ if\ the\ box\ j\ is\ in\ the\ cell\ i;\ 0\ otherwise.$
- $(x, y)$ y $(w, h)$ : are the coordinates, and the height and width of the predicted box.
- $(\hat{x}, \hat{y})$ y $(\hat{w}, \hat{h})$ : are the coordinates, and the height and width of the ground truth bounding box.
- $\lambda_{coord}$ y $\lambda_{noobj}$: Are regularization constants.
- $B$: The number of found boxes.
- $S$ : The number of cells where the image is divided.
- $C$: The confidence score of the presence of the object.
- $\hat{C}$: The IoU coefficient of the predicted bounding box and the ground truth.
- $\rho_i(C)$: The predicted class.
- $\hat{\rho}_i(C)$: The ground truth class.

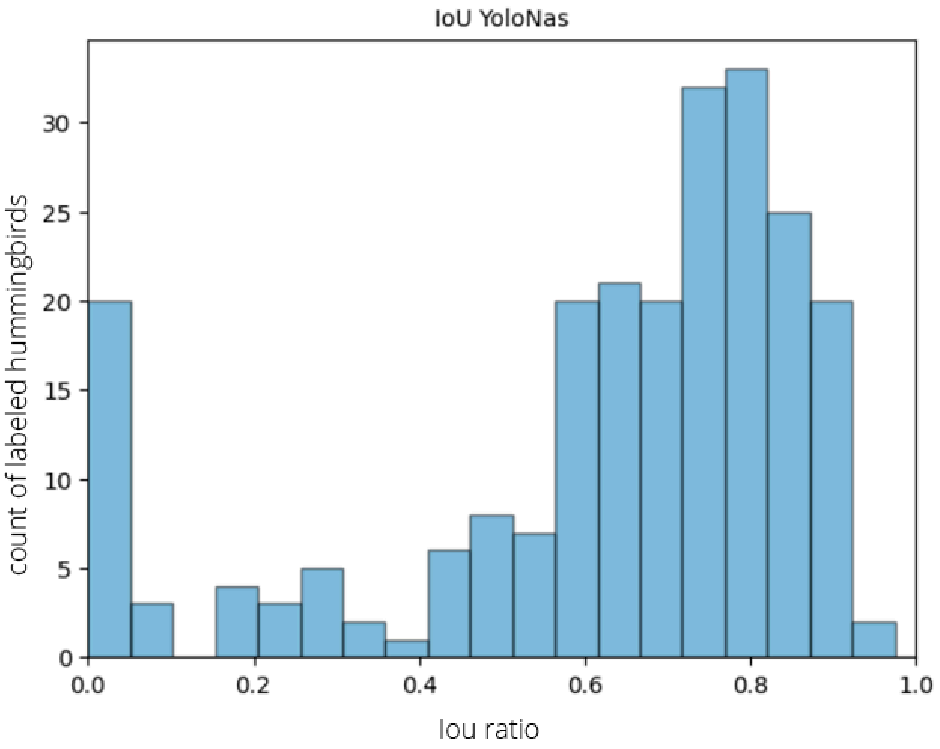


**Figure S1.** Histogram of the iou ratio for the detection process. Here the IoU is obtained from the manual labeled bounding box against the predicted bounding box.

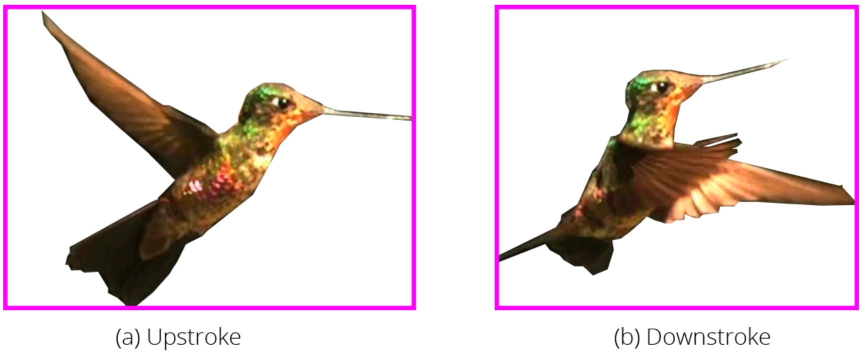


**Figure S2.** Hummingbird wingbeat analysis. (a) Upstroke and (b) Downstroke. The boxes for both actions remain very similar to each other.

**Table S1.** TP, FP, and FN for the classification stage and each analyzed video. We only focused on the class hummingbird, and analyzing the objects detected by the detection stage.

| **Species** | **Sex** | **TP** | **FN** | **FP** | **TN** |
|---|---|---|---|---|---|
| *Saucerottia saucerottei* | U | 1 | 0 | 1 | 1 |

| | | | | | |
|---|---|---|---|---|---|
| *Amazilia tzacatl* | U | 3 | 0 | 0 | 0 |
| *Chalybura buffonii* | M | 1 | 1 | 0 | 0 |
| *Coeligena helianthea* | F | 1 | 0 | 0 | 0 |
| *Colibri coruscans* | F | 1 | 0 | 0 | 1 |
| *Coeligena phalerata* | F | 1 | 0 | 0 | 1 |
| *Eriocnemis cupreoventris* | F | 1 | 0 | 0 | 1 |
| *Ensifera ensifera* | M | 1 | 1 | 0 | 1 |
| *Eriocnemis vestita* | M | 1 | 0 | 0 | 2 |
| *Metallura tyrianthina* | F | 1 | 0 | 0 | 1 |
| *Thalurania colombica (1)* | F | 1 | 0 | 0 | 1 |
| *Thalurania colombica (2)* | F | 1 | 1 | 0 | 1 |
| *Thalurania colombica (3)* | M | 0 | 1 | 0 | 0 |

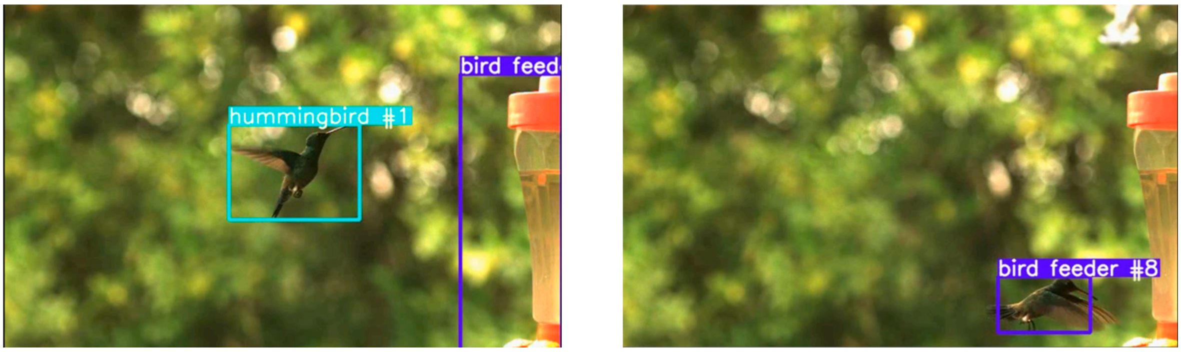


**Figure S2.** Classification example, two frames of the same videos counting as TP and FN. (a) Shows the correct classification of a hummingbird, while (b) shows a misclassification of the same hummingbird in the same video.

**Table S2.** Results obtained by the algorithm in the columns Count of peaks and Frequency algorithm, and its comparison with the ones obtained by the expert biologist: Count wingbeats and Frequency ground truth.

| Species | Final Frame | Count peaks algorithm | Frequency algorithm (Hz) | Count wingbeats manually | Frequency manually (Hz) | Error Freq. (%) |
|---|---|---|---|---|---|---|
| *Saucerottia saucerottei* | 1279 | 75 | 35.08 | 91 | 35.57 | 1.4 |
| *Amazilia tzacatl* | 395 | 32 | 31.13 | 25 | 31.65 | 1.6 |
| *Chalybura buffonii* | 410 | 25 | 32.22 | 23 | 28.05 | 14.9 |
| *Coeligena helianthea* | 442 | 26 | 30.44 | 22 | 24.89 | 22.3 |
| *Colibri coruscans* | 355 | **18** | 25.35 | **18** | 25.35 | **0** |
| *Coeligena phalerata* | 996 | 59 | 30.29 | 67 | 33.63 | 10 |
| *Eriocnemis cupreoventris* | 331 | 18 | 31.36 | 19 | 28.7 | 9.26 |
| *Ensifera ensifera* | 1067 | 31 | 31.76 | 142 | 66.54 | 52.27 |
| *Eriocnemis vestita* | 1012 | 10 | 31.85 | 58 | 28.66 | 11.14 |
| *Methalura tyrianthina* | 305 | 20 | 36.1 | 18 | 29.51 | 22.34 |
| *Thalurania colombica (1)* | 372 | 20 | 29.15 | 28 | 37.63 | 22.53 |
| *Thalurania colombica (2)* | 307 | 10 | 34.48 | 27 | 43.97 | 21.58 |
| *Thalurania colombica (3)* | 362 | 25 | 34.25 | 31 | 42.82 | 20.02 |

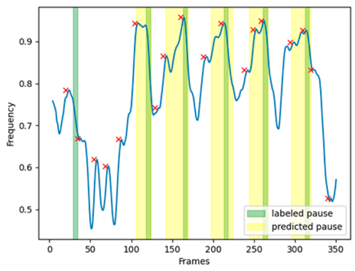


video name: Colibri

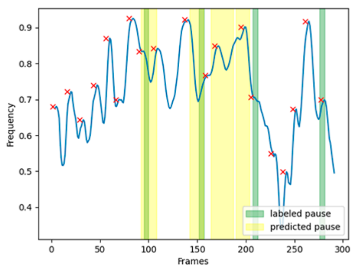


video name: Mtyrianthina

**Figure S3.** Frequency iou signal vs frames plots of two different videos. The signal peaks are found and count for the measure of the final frequency flight.

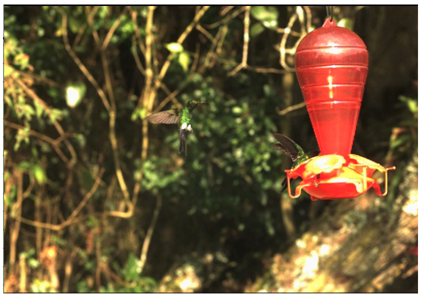

***(a) E. vestita***

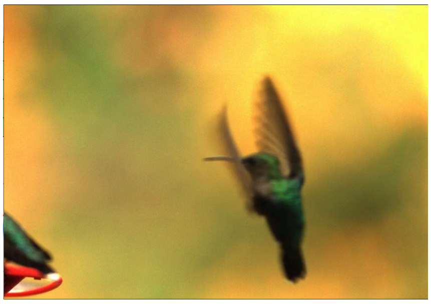

**(b) *T. colombica (3)***

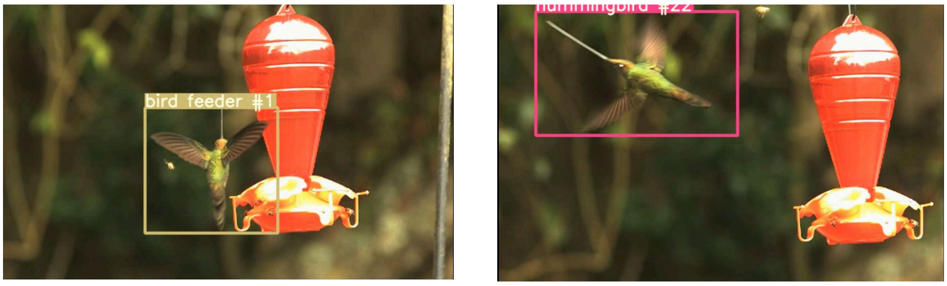


**(c)** ***E. ensifera***

**Figure S4.** Frame shot from the videos where the error of the count was bigger than 50 %. Video names are: *E. vestita* with an error of 70 %, and *T. colombica* (2) with an error of 59 %

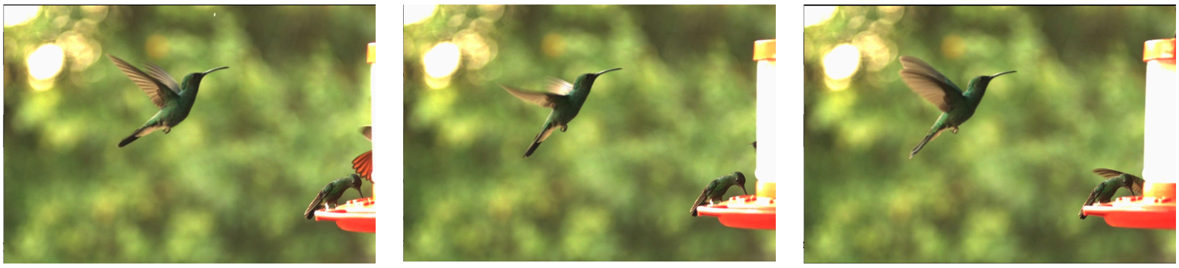

**(a)** ***C. buffonii***

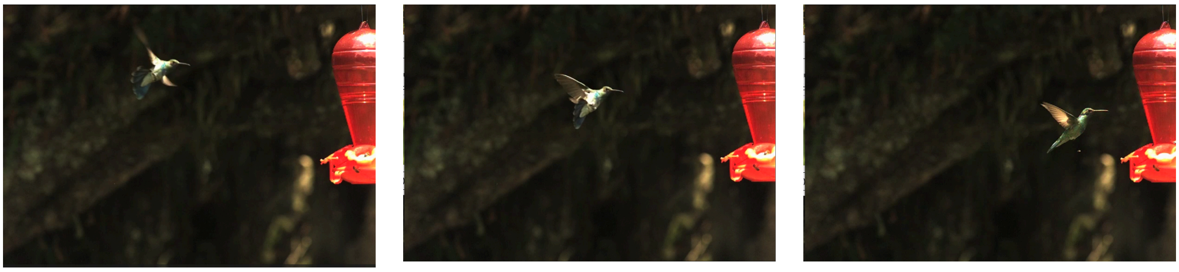

**(b)** ***C. coruscans***

**Figure S5.** Frame shot from the videos where the error of the count was 0 %. Hummingbirds are in the center of the scene and it could be well differentiated from the background

We provide a list of suggestions for improving the filming and implementation of the proposed algorithm (Supplementary Information) for future studies.

1. **Optimal Timing**: Capture videos during early morning or late afternoon when hummingbirds are most active. This also helps in avoiding harsh midday sunlight, which can cause glare and shadows.
2. **Stable Backgrounds**: Choose locations with minimal background movement to reduce distractions in the video. Natural settings with consistent foliage or flowers work well.

3. **Lighting Conditions**: Use natural light to your advantage. Avoid direct sunlight which can create harsh contrasts. Overcast days provide diffused lighting, reducing shadows and highlights.
4. **Consistent illumination:** Abrupt illumination changes could affect the performance of the tracking algorithm.
5. **Camera Setup**: Use high-speed cameras to capture the rapid wing movements. Ensure the camera has a high frame rate (at least 240 frames per second) to accurately capture wingbeat frequency.
6. **Mounting and Stabilization**: Use tripods or fixed mounts to keep the camera steady. This helps in getting clear, stable footage without motion blur.
7. **Proximity and Zoom**: Position the camera close enough to the hummingbird's feeding area but not too close to disturb them. Use optical zoom to get detailed footage without compromising video quality.
8. **Keep an optimal focus:** Blurry frames smooth the high frequency features of the image, losing important details for the system.
9. **Background Color and Contrast**: Ensure the background contrasts well with the hummingbird's colors. A plain, consistent background makes it easier for machine learning algorithms to detect and analyze the bird's movements.
